# Enhanced identification of small molecules binding to hnRNPA1 via cryptic pockets mapping coupled with X-Ray fragment screening

**DOI:** 10.1101/2024.12.17.628909

**Authors:** Louise Dunnett, Sayan Das, Vincenzo Venditti, Filippo Prischi

**Affiliations:** Diamond Light Source Ltd., Harwell Science and Innovation Campus, Didcot OX11 0QX, UK; Department of Chemistry, Iowa State University, Ames, Iowa 50011, United States; Randall Centre for Cell and Molecular Biophysics, King’s College London, London, SE1 1UL, UK

## Abstract

The human heterogeneous nuclear ribonucleoprotein (hnRNP) A1 is a prototypical RNA-binding protein essential in regulating a wide range of post-transcriptional events in cells. As a multifunctional protein with a key role in RNA metabolism, deregulation of its functions has been linked to neurodegenerative diseases, tumour aggressiveness and chemoresistance, which has fuelled efforts to develop novel therapeutics that modulates its RNA binding activities. Here, using a combination of Molecular Dynamics (MD) simulations and graph neural network pockets predictions, we showed that hnRNPA1 N-terminal RNA binding domain (UP1) contains several cryptic pockets capable of binding small molecules. To identify chemical entities for development of potent drug candidates and experimentally validate identified druggable hotspots, we carried out a large fragment screening on UP1 protein crystals. Our screen identified 36 hits which extensively samples UP1 functional regions involved in RNA recognition and binding, as well as mapping hotspots onto novel protein interaction surfaces. We observed a wide range of ligand-induced conformational variation, by stabilisation of dynamic protein regions. Our high-resolution structures, the first of an hnRNP in complex with a fragment or small molecule, provides rapid routes for the rational development of a range of different inhibitors and chemical tools for studying molecular mechanisms of hnRNPA1 mediated splicing regulation.

## Introduction

RNA-binding proteins (RBPs) hold the vital role to regulate post-transcriptional events in every cell. RBPs recognition and binding to RNA sequences and/or secondary structure motifs are highly dynamic and tightly regulated processes. In humans there are about 2,000 RBPs, grouped into several families based on the presence of structurally conserved RNA-binding domains (1,2). The heterogeneous nuclear ribonucleoprotein (hnRNP) A1 is a prototypical RBP implicated in multiple aspects of nucleic acid metabolism, including processing of micro-RNA precursors, transcription regulation, constitutive and alternative splicing, and nucleo-cytoplasmic mRNA transport (3). hnRNPA1 ability to carry out these wide range of functions is encoded in its modular domain organisation consisting of an N-terminal Unwinding Protein 1 (UP1) domain and a C-terminal intrinsically disordered glycine-rich domain (4). UP1 is composed of two RNA Recognition Motifs (RRM1 and RRM2) which, despite having a high level of sequence similarity and recognising similar optimal motifs, 5’-YAG-3’ (Y is a pyrimidine), are non-redundant and functionally non-equivalent (4). Over the last 20 years several structural studies of the protein in complex with telomeric DNA, RNA and miRNA have provided ligand binding models, although the molecular mechanism driving selective DNA/RNA recognition remain elusive (5).

As a major regulator of gene expression, deregulation of hnRNPA1 functions has been linked to several pathological cellular conditions. A growing number of studies has suggested that hnRNPA1 contributes to development of neurodegenerative diseases (6). For example, hnRNPA1 mutations have been identified to alter splicing, often by causing exon-skipping events, and to increase formation of self-seeding fibrils in Multiple Sclerosis (7), and Amyotrophic Lateral Sclerosis (8), respectively. Furthermore, altered hnRNPA1 expression has been found to enhance translation of dynamin-related protein 1 (Drp1) (9), a GTPase protein often overexpressed in Huntington’s disease, and affect the alternative splicing of amyloid precursor protein (APP), causing increased secretion of β-amyloid peptide (10). HnRNPA1 has also been shown to promote viral replication. Specifically, several viruses (e.g., Human Rhinovirus, Enterovirus, Sindbis virus, and Human Immunodeficiency Virus I) exploit hnRNPA1 internal trans activating factor (ITAF) activity, resulting in an increased Internal Ribosome Entry Site (IRES) mediated translation of viral RNA (11–14). Deregulated hnRNPA1 expression or activation via posttranslational modifications has been shown to increase cell proliferation and survival in various cancer types, including lung, breast, prostate and gastric cancers (15–19), leukemia (20), Burkitt lymphoma (21), multiple myeloma (22), hepatocellular and cervical carcinomas (23,24). Although hnRNPA1 has been shown to be involved in a wide range of molecular events driving tumorigenesis and drug resistance, the underlaying molecular mechanism seems to be linked to an alternation in RNA recognition and binding, leading to an increased translation of pro-survival proteins and oncoprotein variants (3). Consistently, it has been shown that knockdown of hnRNPA1 leads to apoptosis preferentially in cancer cells over their healthy counterparts (25).

Due to the direct and indirect involvement of hnRNPA1 in neurodegenerative disorders, viral gene expression, and carcinogenesis, preventing or fine-tuning hnRNPA1-RNA binding with novel therapeutics is an emerging area of scientific interest, especially considering the growing number of RBPs inhibitors showing promising pre-clinical activities (26). Earlier attempts to target hnRNPA1 identified a small number of compounds that alter the protein activity in different ways. For example, Carabet et al. (27) showed that the small molecule VPC-80051 binds to the RRM1 of hnRNPA and may act as a RNA competitor, causing a small reduction of the Androgen Receptor splice variant (27). Quercetin, a promiscuous naturally occurring flavonoid, has been shown to bind the unstructured C-terminal region of hnRNPA1, preventing hnRNPA1 nucleocytoplasmic shuttling and causing cytoplasmic accumulation (28). Similarly, Camptothecin, a non-competitive protein-protein interaction inhibitor, binds the hnRNPA1 C-terminal and prevents its interaction with Topoisomerase I (29). Although promising, these compounds show limited selectivity or specificity for hnRNPA1. A major obstacle in the identification of suitable inhibitors is the lack of easily identifiable deep binding pockets. Here, to overcome this limitation and identify small molecules ligands of hnRNPA1, we used Molecular Dynamics (MD) simulations and graph neural network pockets predictions coupled with X-Ray fragment screening. Our simulations showed that hnRNPA1 N-terminal RNA binding domain (UP1) contains several cryptic pockets (i.e., concavities, often absent in crystal structures, that open on the protein surface when the protein fluctuates to an excited state) capable of binding small molecules. Using the XChem facility at the Diamond synchrotron, we carried out a large fragment screening on UP1 protein crystals. Our screen identified 36 hits which extensively samples UP1 functional regions involved in RNA recognition and binding, as well as mapping hotspots (i.e., a cluster of residues that makes a major contribution to the binding free energy) onto novel protein interaction surfaces. Our high-resolution structures, the first of an hnRNP in complex with a fragment or small molecule, provides rapid routes for the rational development of a range of different potent inhibitors and chemical tools for studying molecular mechanisms of hnRNPA1 mediated splicing regulation.

## Materials and Methods

### Molecular Dynamics simulations and hot spot mapping

A 2 μs simulation was performed by using Gromacs 2023 package (30) with Amber03 force field (31). The NMR solution structure (PDB ID: 2LYV) (32) was used as a starting conformation. The system was placed in a TIP3P water box with a 1 nm distance between the boundary of the box and the system. One Cl^-^ counterion was added to neutralize the charge. Energy minimization was carried out using the steepest descent method with a step size of 0.01 nm. Minimization was run until the maximum force fell below 1000 kJ/mol/nm for 50000 steps. The system equilibrated at 300 K for 100 ps with a timestep of 2 fs for 50000 steps and equilibrated at a constant pressure (1 atm) for 100 ps with a timestep of 2 fs for 50000 steps, both using the leap frog integrator. LINCS was used to constrain the covalent hydrogen bonds with a Verlet cutoff scheme with a 1.0 nm cutoff radius for neighbour list. The particle mesh Ewald scheme was used to treat long range interaction with a Fourier grid spacing of 0.16 nm. The modified Berendsen thermostat (velocity-rescale thermostat) and the Parrinello-Rahman barostat were used to maintain the temperatures at 300 K and 1 atm. To generate an ensemble of structures, the MD trajectory was clustered by k-means clustering using MDAnalysis (33). 50 clusters were generated, and the representative structure of each cluster are available here https://github.com/fprischi/UP1_clusters. Cryptic pockets were identified by analysing the MD trajectory using fpocket 2.0 (34).

The NMR ensemble (PDB ID: 2LYV) was analysed using PocketMiner (35) and we consider only pockets with a likelihood (averaged for the ensemble) greater than 0.5.

### UP1 preparation for structural studies

The N-terminal (His)_6_UP1 fusion was expressed and purified as previously described (32). In short, BL21(DE3) (Invitrogen) were transformed with a pETM-14 vector containing the DNA sequence encoding the UP1 domain of hnRNP A1 (residues 2–196) (Uniprot entry P09651). Cells were grown in LB media overnight at 18 °C, resuspended in lysis buffer (50 mM Tris-HCl, pH 8.00, 300 mM NaCl, 0.01 % (v/v) Triton X-100, 20 mM imidazole, 5 % (v/v) glycerol, 5 mg/mL lysozyme from hen’s egg white, 1.0 mg/mL DNase from bovine pancreas, EDTA-free protease inhibitor (Mini Tablets, Pearce)) and lysed by sonication. Cell lysate was centrifuged at 18,000 x g for 1 h at 4 °C, supernatant was loaded on a Ni–NTA column (Cytiva), and (His)_6_UP1 was eluted with an imidazole gradient. The (His)_6_ tag was removed via digestion overnight with Human Rhinovirus 3C protease (PreScission protease) at 4 units/mL at 4 °C. UP1 was then loaded on a HiTrap HP SP column (Cytiva) equilibrated with 20 mM MES pH 6.00, 5 mM β-mercaptoethanol, 1 mM EDTA, 5 % (v/v) glycerol, and was eluted with an NaCl gradient. Fractions containing UP1 were pooled and loaded onto a HiLoad 16/60 Superdex 75 (Cytiva) equilibrated with crystallisation buffer (20 mM MES pH 6.00, 150 mM NaCl, 5 mM β-mercaptoethanol, 1 mM EDTA, 5 % (v/v) glycerol).

### UP1 Sitting Drop Crystallisation

Protein from a single purification batch was used for in crystallo fragment screening in 96-well MRC 2 lens crystallisation trays (SWISS-Sci). All reagents were pre-equilibrated to 16 °C prior to use. A liquid handling robot (Mosquito, TTP Labetch) was used to dispense UP1 (from a 15 mg/mL stock) and precipitant solution (0.1 M Tris-HCl pH 8.50, 25% (w/v) PEG-4000 and 8% (v/v) MPD) as 400 nl sitting droplets at a 1:3 ratio of protein:precipitant. Each plate was immediately incubated at 16 °C following protein dispensing.

### Droplet Targeting using TexRank and Fragment Soaking

Soaking, mounting, and data collections were performed following the XChem facility workflow (36). Crystal droplets were imaged using a Rock Imager (Formulatrix), and targeted for fragment droplet dispensing using TexRank software (37). A total of 768 fragments from the DSI poised library (38) (500 mM stock concentration in DMSO) were dispensed into the crystal droplets using ultrasonic sound waves by a liquid handling robot (ECHO 550, Labcyte, Inc.) (39), with a 100 mM final concentration of fragment (20% (v/v) final concentration DMSO) in the droplets. Crystals were incubated at 16 °C for 3 hours before being mounted with the aid of a Crystal Shifter (Oxford Lab Technologies) and flash frozen in liquid nitrogen. Automated data collection was performed on the I04-1 beamline with a Pilatus 6M-F detector (Dectris, Switzerland) using a fixed wavelength of 0.92 Å and a beam size of 60 x 50 mm on cryo-cooled crystals.

### Structure solution and refinement

The diffraction images were processed using *xia2 3dii* or *xia2 dials* (40,41), followed by AIMLESS (42). UP1 crystals belonged to space group P 1 21 1 and consisted of one protein per asymmetric unit. After molecular replacement and refinement of the initial model, the resulting maps were analysed by PanDDA (43) followed by model building using Coot 0.9.8.93 (44). The structures of confirmed hits were then refined using PHENIX 1.20.1 (45).

## Results

### Computational hotspot mapping

MD derived structures have been extensively used to sample small conformational rearrangements and map hotspots and cryptic pockets on protein surfaces, showing good agreement between predicted and experimentally mapped pockets (46,47). We focused our studies on UP1 domain, as hnRNPA1 C-terminal is intrinsically disordered and not suitable for structural studies. Differently, the two RRMs in UP1 have a conserved βαββαβ structure with a four-stranded antiparallel β-sheet sitting over two helices, joined together by highly dynamic inter-RRMs loop (Fig. 1A) (4). To identify cryptic pockets on UP1 surface we performed a 2 μs MD simulations, which allows to take into account the effects protein conformational dynamics in a water environment have in the formation of binding cavities. In line with previous similar studies (48), we observed good overall agreement between the root mean square fluctuation (RMSF) derived from our MD simulations and the RMSF calculated using the experimentally determined UP1 NMR structural ensemble (PDB ID: 2LYV) (Fig. S1). We then generated an ensemble of structures using the MD trajectories, which were used to map and characterise cryptic pockets using fpocket 2.0 (34). To further validated identified pockets, we compared fpocket results with pockets predicted using PocketMiner (35). This robust approach increased reliability that the predicted pockets exist on UP1 and that these pockets are also suitable for small molecule binding.

**Figure 1.**
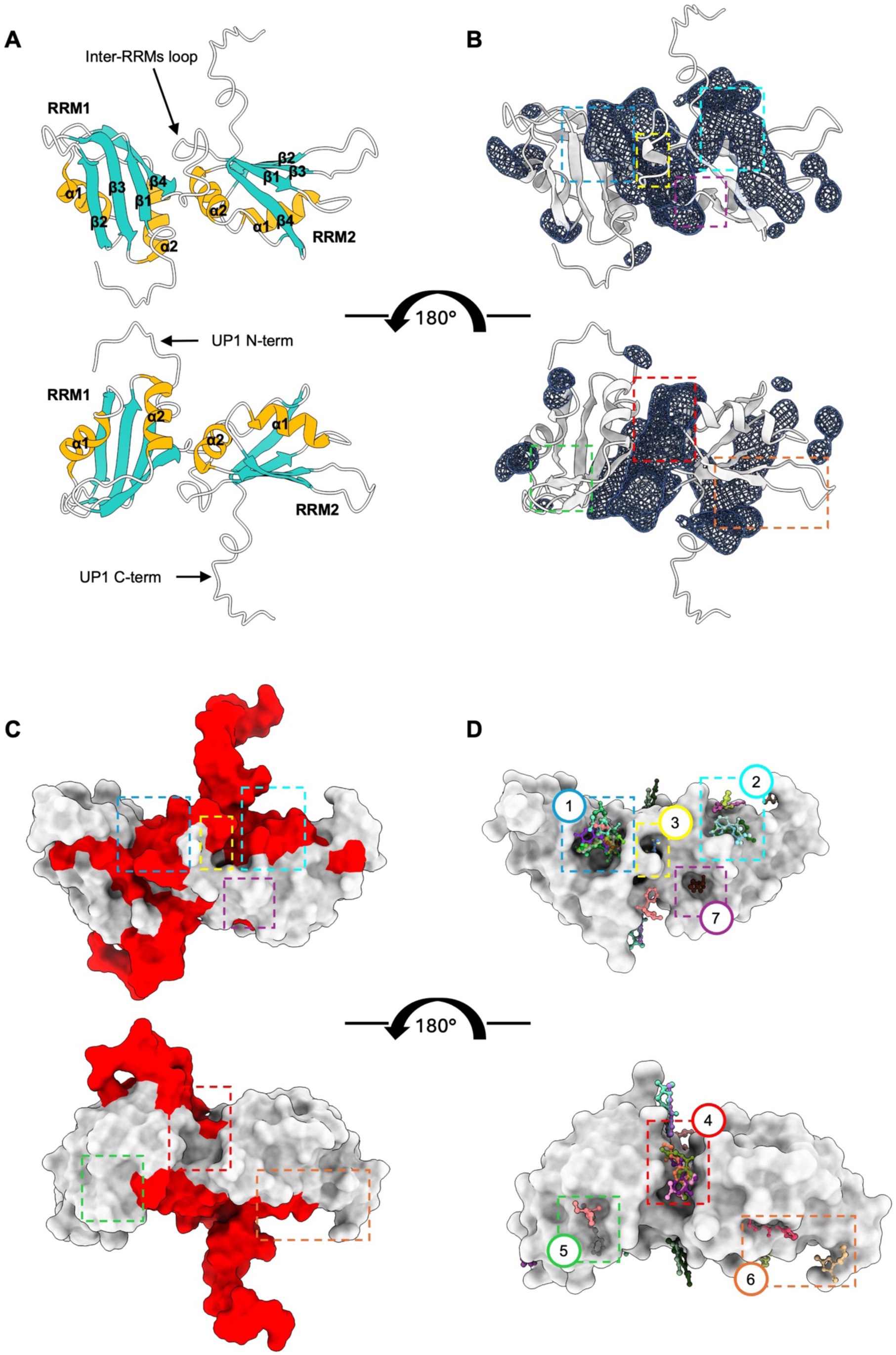
UP1 cryptic pockets identification. **A**) Cartoon representation of UP1, with labelled conserved secondary structures. Flexible regions (N- and C-termini and the inter-RRMs loop) are visible in the NMR structure but not in most X-Ray structure of the apo UP1. **B**) UP1 cartoon representation (grey) with cryptic pockets identified via MD simulations, shown as blue isosurfaces. **C**) UP1 surface coloured according to PocketMiner predicted cryptic pocket likelihood, >0.5 red and <0.5 grey. **D**) UP1 crystal structure with all fragment hits shown as ball-and-sticks, and druggable hotspots highlighted by coloured boxes. For reference, coloured boxes in (D) are also shown on (B) and (C). **1**: nucleobase pocket 1; **2**: nucleobase pocket 2; **3**: inter-RRMs loop (hidden behind the loop in (A) and (B)); **4**: inter-RRMs (ɑ2-ɑ2); **5**: RRM1 ɑ1 - (ɑ2-β4) loop; **6**: RRM2 ɑ1 - β2 and the adjacent RRM2 (β2-β3) loop sites; **7**: RRM2 ɑ2 - β4. All structures shown have the same orientation.

Both approaches identified similar pockets located on UP1 functional regions. Specifically, two hotspots were identified on the two UP1 binding platforms or nucleobase pockets (49), created by the β-sheets of the RRM1 and the inter-RRMs loop (RRM1 nucleobase pocket), and the β-sheets of the RRM2 and the UP1 C-terminal (RRM2 nucleobase pocket), thus highlighting the key role the highly dynamic inter-RRMs loop and the UP1 C-terminal have in the formation of these binding surfaces (Fig. 1B-C). A small difference is visible for the RRM2, where PocketMiner suggests that the region in between β1-β3, identified in our MD simulation as part of a large cryptic pocket on the RRM2 nucleobase pocket, is not a likely pocket (Fig. 1C). The largest hotspot region predicted by the MD simulation extends throughout the whole RRM1-RRM2 interface (or inter-RRMs interface or inter-RRMs (ɑ2-ɑ2)) and terminates at the base of the UP1 C-terminal tail, while PocketMiner suggests that only part of this surface is viable for small molecules binding (Fig. 1B-C). This inter-RRMs interface, stabilised via two salt-bridges between R88-D157 and R75-D155 and a hydrophobic cluster involving L13, I164 and V90, is key for protein function (32,50). Indeed, recent studies have shown that the inter-domains interactions in the inter-RRMs interface have a role in allosterically regulating hnRNPA1 (5), and that the allosteric coupling of RRM1 and 2 allows fine-tuning of hnRNPA1’s affinity for different RNA with optimal or near optimal consensus motifs (4,49,50). Finally, smaller binding pockets are visible at the edges of the two RRMs, likely generated by dynamics of the flexible RRMs β2-β3 loops (Fig. 1B), none, however, detected by PocketMiner (Fig. 1C).

Taken together these data suggest that several druggable cryptic pockets exist on the hnRNPA1 N-terminal RNA binding domain surface, which encouraged us to carry out a crystallographic fragment screening.

### Large fragment screening on UP1 protein crystals

There is currently no experimental structural information for hnRNPA1, or any other member of the hnRNP family, in complex with small molecules or drug-like compounds. To overcome this limitation, we carried out high-throughput fragment screening on protein crystals, a relatively novel and powerful approach to identify chemical entities for development of potent drug candidates and chemically probe protein surface for the identification of druggable hotspots (51,52). We screened the Diamond-SGC-iNext Poised Library, comprising 768 chemically diverse fragments (38,53), directly on UP1 crystals. Out of 548 successfully collected datasets, PanDDA (43) identified 61 potential hits, which were manually verified in Coot (44), resulting in 36 refined high resolution UP1 fragment-bound structures (Table S1). Nearly 80% of all the identified hits occupies known functional regions of UP1 and computationally predicted bind sites (Fig. 1D), with 48% and 13% identified fragments binding to the RRM1 and 2 nucleobase pockets, respectively, 13% on the inter-RRMs (ɑ2-ɑ2), and 6% on the inter-RRMs loop (Fig. 1D). The high level of agreement between experimentally identified hotspots and predicted cryptic pockets increases reliability of identified hits and released models, and improves accuracy in the prioritization of hits. The other hits are found onto a likely conserved RRM protein-protein interaction (PPI) surface (Table S3). The majority of compounds showed high binding surface selectivity, and only seven bind at the interface of two or three symmetry-related molecules, two of which were also identified on two different binding surfaces (Table S3).

### Ligand-induced conformational variations on the RRM1

Fifteen fragments were identified to bind on the RRM1 nucleobase pocket, all interacting with key residues in the two ribonucleoprotein consensus sequences (RNPs). Indeed, extensive structural studies investigating UP1 mode of binding to the consensus motifs 5’-YAGG-3’ have shown that RNPs mediate sequence specific ssRNA binding. Specifically, the conserved phenylalanines in the RNPs (F17, F57, F59) directly interact with the central A_2_G_3_ nucleotides, with the central A_2_ stacking in between F17 and H101, the G_3_ stacking onto F59 and forming van der Waals contacts with F57, and the phosphate group between A_2_ and G_3_ forming an electrostatic interaction with R55 (50,54). Additional residues in the RRM1 nucleobase pocket, E85/K87 and D42/R92, form a network of hydrogen bonds with the flanking Y_1_ and G_4_, respectively (49,50,54). Interestingly, all fragments identified on the RRM1 nucleobase pocket formed van der Waals interactions with the conserved F17, F57 and/or F59, and frequently π-stacking with F17 (Table S4) through aromatic ring or similar heterocycle. Importantly, we observed a series of highly relevant conformational variations in the sidechains of the key conserved F17 and F59 in the RNPs of RRM1 (Table S3). Earlier studies by Vitali et al. (55) showed that in UP1 apo structure F17 and F59 have correlated alternative conformations, with side chains adopting (i) F17A-F59A orientation (RNA-bound conformation), seen also in the UP1-DNA and -RNA bound structures and the only conformation seen in other RRM-nucleic acid complex structures (49,54); and (ii) F17B-F59B (anti-RNA-bound conformation), not frequently seen in other RRM structures, including the RRM2 of UP1. Fragments binding to the RRM1 nucleobase pocket modulate these dynamics feature by recognising different RRM1 RNPs conformations (Table S3). Three fragments (Z1373445602, Z1401276297, Z106579662) sit on the RRM1 β-sheet in an orientation that closely resemble that of purine ring of the central A_2_, with their aromatic rings sandwiched between F17 and H101, stabilising the F17 side chain in an RNA-bound conformation but not altering F59 dynamics (Fig. 2A). We also observed the opposite conformational change, likely induced by the presence in the fragments (ZINC72259689, Z1152242726, Z137811222, and Z1217960891) of a large cycle at one end of the fragment positioned in proximity of the F59, which locks its side chain in an anti-RNA-bound conformation, and a nitrogen containing cycle at the other end, which interacts with either conformation of the F17 (Fig. 2B). Three fragments are positioned parallel to the RRM1 β-sheet direction, in an orientation similar to that occupied by the sugar-phosphate backbone in the UP1-DNA bound structures (49,54). These three fragments (Z56880342, Z54508609, Z1220452176) have a similar geometry and groups composition, with an aromatic head stacked in the middle of F17, F57 and F59 and a polar tail protruding outward. This induces a conformational variation of the side chains of F17 and F59 which adopt (mainly or only) an anti-RNA-bound conformation (Fig. 2C) (49,54). Only four fragments reach the inter-RRMs loop and alter the RRM1 nucleobase pocket structure. (i) ZINC72259689 and Z1152242726 form H-bonds with the V90 backbone, partially rigidifying the inter-RRMs loop and pushing the H101 sidechains backward (toward the RRM2) of 7.5 Å compared to the Cγ relative position in the DNA bound structure (Fig. 2B) (residue not visible in the apo structure). Differently (ii) Z991506900 and Z30820160 coordinate several water molecules, with a much higher number of visible water molecules (not present in other structures) bridging the inter-RRMs loop. This is likely linked to a reduction in flexibility of the loop (fully visible in both structures), which folds back into the RRM1 nucleobase pocket, creating a pronounced positive charged binding cavity in which the fragments are lodged (Fig. 2D and S1A). Interestingly, this also induce an RNA-bound conformation for both F17 and F59 side chains (Table S3).

**Figure 2.**
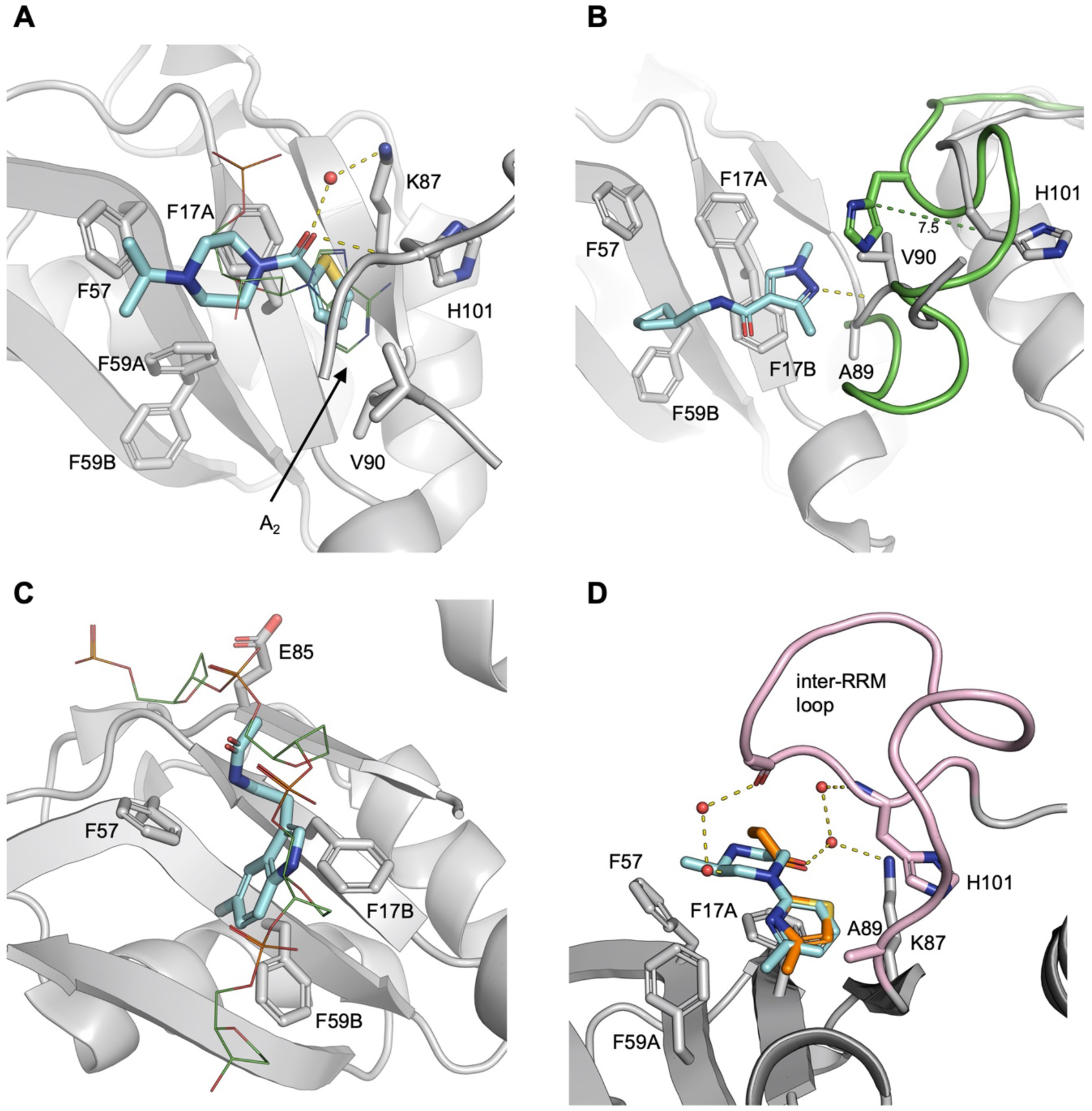
RRM1 nucleobase pocket. **A**) Overlay of Z106579662 (cyan sticks) and A_2_ (PDB ID: 2UP1) (green lines). Residues mediating interaction are shown as grey sticks, highlighting F59 alternative conformations and F17 RNA bound conformation. **B**) ZINC72259689 (cyan sticks) stabilizes F59 in anti-RNA-bound conformation but doesn’t affect F17 alternative conformations. The UP1-ZINC72259689 structure (grey cartoon) is overlayed to UP1-DNA structure (PDB ID: 2UP1, with inter-RRMs loop shown as green cartoon). The dotted green line highlights the different position of H101 side chain in the two structures. **C**) Overlay of Z1220452176 (cyan sticks) and the DNA sugar-phosphate backbone (PDB ID: 2UP1) (green lines), highlighting F17 and F59 anti-RNA bound conformations. **D**) Overlay of Z30820160 (orange sticks) and Z991506900 (cyan sticks), with F17 and F59 in RNA-bound conformations. The fully visible inter-RRMs loop is shown in pink. In all panels, water molecules are shown as red spheres, sticks are coloured by heteroatom, and hydrogen bonds with lengths 2.5Å-3.5Å are shown as yellow dotted lines.

### Fragment hits reveal multiple targetable hot spots

Similar to RRM1, fragments binding to the RRM2 nucleobase pocket interacts with the hydrophobic residues on the RNPs (F108, F148) (Fig. 3A), occupying a position similar to that of purine rings of either the A_2_ (Z416341642, Z992569480, Z906021418) or G_3_ (Z641230552, Z641239276).

**Figure 3.**
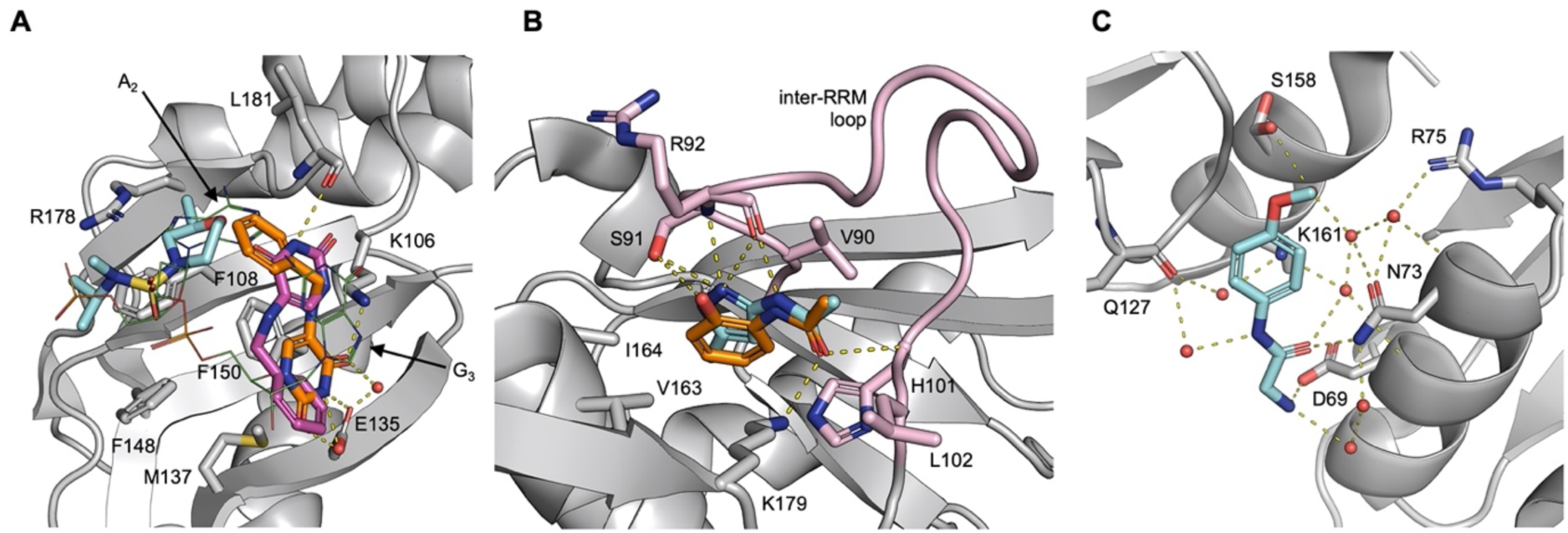
Fragments extensively sample UP1 functional regions. **A**) Overlay of Z416341642 (cyan sticks), Z641230552 (with the two alternative locations in orange and purple) and DNA (PDB ID: 2UP1) (green lines) bound to the nucleobase pocket 2. **B)** Overlay of Z57040482 (orange sticks) and Z237527902 (cyan sticks) bound to the inter-RRMs loop site. The fully visible inter-RRMs loop is shown in pink. **C)** Inter-RRMs (ɑ2-ɑ2) site with EN300-805013 (cyan sticks). In all panels, sticks are coloured by heteroatom, water molecules are shown as red spheres, and hydrogen bonds with lengths 2.5Å-3.5Å are shown as yellow dotted lines.

Differently, binding of fragments on the inter-RRMs loop and the inter-RRMs (ɑ2-ɑ2) is mostly driven by polar interactions. Two fragments (Z57040482, Z237527902) bind directly the inter-RRMs loop. This induces a conformational variation causing the loop to fold toward RRM2 ɑ2β4 and form a deep pocket (Fig. S2B), similar to the one present in the UP1-RNA/DNA bound structures (49,54), but absent in the apo structure (55). These fragments form H-bonds between the backbone amino group of L102 backbone, the side chain of K179 and a carbonyl group in an acetamide moiety, and between the backbone amino group of R92 and the side chain of S91 and the phenyl group in Z57040482 and the nitrogen containing heterocycle of Z237527902 (Fig. 3B, Table S4). The four fragments identified in the inter-RRMs (ɑ2-ɑ2) contain one or two amide or sulfonamide group aligned parallel to the inter-RRMs interface and forming a series of hydrogen bonds or water bridges with several residues from both the RRM1 (i.e., D69, N73, R75) and the RRM2 (i.e., E126, Q127, K161) (Fig. 3C, Table S4). These hits have expansion vectors suitable for targeting the allosteric pocket containing the inter-RRMs salt-bridges (Fig. S3).

### RRM protein-protein interaction (PPI) surface binders

Earlier studies have shown that hnRNPA1 interacts with several macromolecules, yet there are no experimental structural data on hnRNPA1 in complex with proteins or peptides. However, previous structural studies on other RRM proteins (i.e. FIR, PTBP1, hnRNPL, RBM7, CBP20) (56–61) have shown that the surface defined by α1 - α2 – (α2-β4) loop is a Protein-Protein Interaction (PPI) surface. In all structures, a conserved hydrophobic depression on the PPI surface drives protein/peptide binding via hydrophobic interaction. This hydrophobic depression is also present on both RRM domains of UP1 (Fig. S4). Considering the RRM fold conservation, it is tempting to speculate that these regions are also key in mediating PPI in UP1. While PPI surfaces lack the classic small molecules binding site features detected by MD hotspot mapping and PocketMiner, several small molecules targeting PPI surfaces have been developed to interfere/inhibit interactions (62,63). Importantly, this would recapitulate the presence of five fragments on the RRM1 (Z45617795 and EN300-115958) and RRM2 (Z86417414 and Z802821712, and Z734147462) α1 - α2 – (α2-β4) loop PPI surfaces. (i) Z45617795 and EN300-115958 contain a methanesulfonamide group that forms H-bond with K78, and the phenyl group of Z45617795 is further stabilised by van der Waals interactions with P76 and V83 (Fig. 4A). (ii) The amide group of Z802821712 form H-bond with E118 and I134, while R22 forms H-bond with nitrogen atoms in the pyridazine ring and halogen bond with the fluorobenzene ring of Z86417414 and Z802821712, respectively (Fig. 4B), thus providing an accessible growth vector for targeting the RRM2 PPI surface. Z734147462 forms H-bond between the oxygen atoms of the tetrahydropyran-methanol group and Q165 and V177 (Fig. 4C). Interestingly, this PPI surface is present also in non-canonical RRM domains, which, for example, has been exploited for the design of cyclic peptide and small molecule splicing inhibitors targeting the splicing factor SPF45 (64,65).

**Figure 4.**
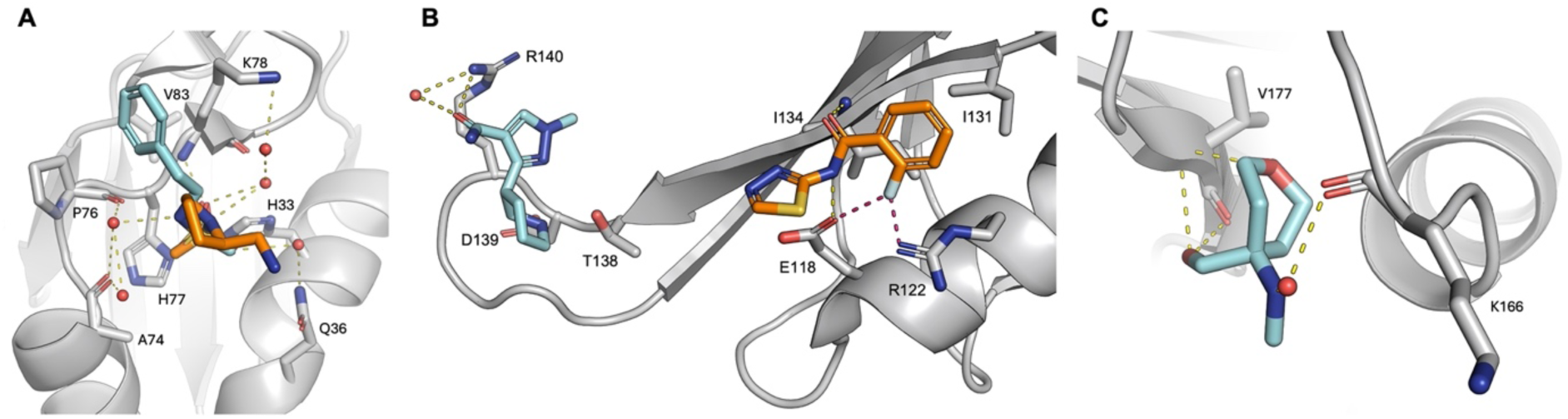
RRM protein-protein interaction (PPI) surface binders. **A**) Overlay of Z45617795 (cyan sticks) and EN300-115958 (orange sticks) bound to the RRM1 ɑ1 - (ɑ2-β4) loop site. **B)** RRM2 ɑ1 - β2 site with Z86417414 (orange sticks) and the adjacent RRM2 (β2-β3) loop site with EN300-197154 (cyan sticks). Halogen Bonds (2.3-2.8Å) are visualised as pink dotted lines. **C**) RRM2 ɑ2 - β4 site with Z734147462 (cyan sticks). In all panels, sticks are coloured by heteroatom, water molecules are shown as red spheres, and hydrogen bonds with lengths 2.5Å-3.5Å are shown as yellow dotted lines.

### Fragments Interactions Influenced by Crystal Contacts

Søndergaard et al. (66) showed that about a third of ligand-bound structures in the PDB have ligand positioned at less than 5 Å distance to a symmetry related atoms and their binding is influenced by crystal contacts and packing. We performed a stringent assessment of the models and identified 7 fragments stabilised by interactions with residues from symmetry related molecules. Of these, six fragments (Z1491353358, Z235361315, EN300-118084, Z198195770, Z104584152, Z106579662) are primarily bound to the inter-RRMs (ɑ2-ɑ2) and additionally stabilised by interaction with residues in the (β2 – β3) loop (Fig. S5A-B), while one (Z641239276) is bound to the RRM2 nucleobase pocket and is additionally stabilised by interaction with residues in the RRM2 α1 – β3 (Fig. S5C). Of particular interest is Z1491353358 which allosterically induce RRM1 RNPs conformational variations (Fig. S2A). Specifically, Z1491353358 induce an RNA-bound conformation for F17 and F59 side chains, potentially as a result of its close proximity to the inter-RRMs loop which shift (1.6 Å and 2.5 Å compared to the A89 relative position in UP1 apo and DNA bound structure, respectively) towards the RRM1 RNPs. This results in an increase of molecular crowding that may force the F17 and F59 side chains into the RNA-bound conformation.

## Discussions

Despite advances in protein-RNA inhibition via targeting of the RNA (i.e., antisense oligonucleotides and short interfering RNAs), RBPs targeting studies are scares, mostly due to the lack of easily identifiable pockets suitable for binding drug-like molecules (67). Most small molecule inhibitors targeting RBPs were identified using standard high-throughput screening of large compound libraries (67). However, an earlier study on HCV NS5b RNA-dependent RNA polymerase showed that fragments with millimolar affinities identified using X-Ray fragment screening could be developed into RBPs inhibitors with nanomolar affinity using structure-based design (68). No similar studies have been carried out on other RBPs, and only a handful of similar X-Ray screening studies have been carried out on DNA binding proteins (69,70). In this paper, we designed a reliable pipeline to perform crystallography-based fragment screen studies on RBPs and tested it on the N-terminal RNA binding domain of hnRNPA1 (UP1), a prototypical RBP. We analysed UP1 protein surface using a combination of MD simulations and graph neural network predictions, and our simulations highlighted the presence of several different cryptic pockets with ideal features for small molecules interaction. To identify compounds able to bind these pockets, we carried out a high-throughput fragment screening on protein crystals, and we solved the first experimental structures of hnRNPA1 (also the first structure of a member of the hnRNP family and an RRM) in complex with small molecules. The bound fragments interact with key protein surfaces, including RNA binding and allosteric sites, and likely PPI surfaces.

It has been shown that proteins are able to assume holo-like conformations even in the absence of interacting ligands (47). Thus, cryptic pockets are present in the apo structures of proteins, and open and become accessible due to protein dynamics. The ligand is then able to recognise its target by “selecting” the most complementary conformation from an ensemble of metastable states and/or by “induced fit”, in either case causing a population shift toward holo states (47). A growing number of studies have shown that these holo-like states can be identified using MD simulations (71,72). Here we performed long MD simulations (2 μs) to identify UP1 holo-like states. While unlikely, even long simulations may fail to sample high-in-energy pockets (73). As such, to robustly and reliably identify cryptic pockets on UP1 we compared pockets identified using MD simulations with pockets predicted using PocketMiner (35). The high level of agreement between predicted pockets strongly suggests that there are several druggable sites on UP1 surface and encouraged us to carry out an X-Ray fragment screening. Indeed, the current lack of experimental structural information is impairing the design and/or identification of compounds with high selectivity or affinity for hnRNPA1.

Small molecules are the most convenient rout to target RBPs, as they present advantageous features, including good solubility and oral bioavailability, ability to cross cell membranes and reach the different compartments where RBPs are localised (74). Despite the lack of experimental structural information, several small molecule inhibitors able to bind to RRM domains and modulate RBPs function have been developed (2). These include compound 27 (and derivatives 838 and 841), which binds directly to the RRM2 of UGBP Elav-like family member 1 (CELF1) and compete with IFN-γ mRNA (75). This prevents mRNA degradation and inhibits hepatic stellate cell activation, suggesting it could be a viable treatment of liver fibrosis (75). Earlier studies identified MS-444 as an inhibitor of Hu antigen R (HuR) dimerisation, probably via binding to its RRM1/2 domains, thus interfering with RNA binding and HuR cellular localisation (76). With a slightly different mechanism, compounds 1c (and derivatives) was predicted to bind in between RRM1 and RRM2 of HuR and compete with RNA binding, showing to potentiate the antitumor effect of standard chemotherapy treatment in breast cancer xenografts (77). Similarly, the small molecule Ro binds the RRM1 of MSI2 and compete with RNA binding, reducing survival of human AML cell lines without affecting normal cells (78). Ro was shown to be selective and not bind other RBPs (78). Indeed, the similarity among all published RRM inhibitors is low (Tanimoto coefficient <0.25 – data not shown) (79), which would recapitulate the observed selectivity. Interestingly, also the similarity among available RRM inhibitors and our identified fragments is similarly low (Tanimoto coefficient <0.25), which would suggest that compounds evolved from fragments described here are likely to be highly selective for hnRNPA1. Similarly to these RRM-inhibitors, 61% of fragments we identified binds directly on the RNA binding surfaces of UP1, suggesting these are likely valuable building blocks for the development of RNA competitors. Furthermore, the ligand induced conformational variation on the RRM1 RNPs we identified may provide different avenues to probe RNA binding. Indeed, recent studies on the RRM domain of CUG-BP2 have shown that RNPs aromatic side-chains motions are critical for RNA recognition (80). The authors also highlighted that these local dynamics are influenced by the conformation of neighbour residues and the binding of RNA is essential to stabilise one uniform set of conformations (80). This is consistent with what we have observed in our structures, where the binding of fragments on the RRM1 nucleobase pocket directly locked F17 and F59 in different sets of conformations. More studies are warranted to better understand how these different conformations can be exploited for small molecule inhibitors targeting. While several studies have successfully identified direct RNA-binding competitors, only a handful of allosteric RBPs inhibitors have been developed. Clingman et al. (81) showed that the binding of ω-9 fatty acids in between the α1-α2 of the RRM of MSI1 induce a conformational change that allosterically inhibits RNA binding. Here we showed that 21% of the identified fragments binds UP1 on regulatory regions outside the RNA binding sites. Of particular interest are fragments binding onto the inter-RRMs interface, as recent studies have shown that this surface is key in relaying thermodynamic stability across UP1 surface, and mutations disrupting RRM1-RRM2 coupling reduce RNA binding and likely hnRNPA1 ability to form dimers (5). Using NMR and MD simulations, a series of key contacts were identified among D69, N73 and R75, and K161, D155, H156 and D157 (5). Compounds Z235361315 and Z33546965 directly interacts with these residues and prevents formation of inter-and intra-RRMs interactions among N73, D157 and K161, providing potential routes for the development of allosteric inhibitors.

Very few papers discuss ligand binding sites located on symmetry-related crystal-packing interfaces, as these are discarded as artifacts and often escapes analysis likely due to (i) not being biological relevant binding sites, potentially the result of high concentration of ligands in the soaking experiment (82), or (ii) having detrimental effects by disrupting contacts between symmetry-related molecules. The few exceptions to this are ligand binding on crystallographic oligomer-interface(s) (83), and molecular glue (84). While the seven ligands we identified on symmetry-related crystal-packing interfaces are unlikely to be viable starting point for further development, they bind in very close proximity to other fragments in the inter-RRMs interface, thus could provide information for chemical merging or linking.

The ability to modulate hnRNPA1 activity in cell of most compounds was assessed (data not shown), but, likely due to the low mM binding affinity and the low solubility, none showed biological activity, in line with what reported in other similar fragment screening studies (68,85). However, the fragments we identified sample extensively the protein surface, identifying novel hot spots and revealing diverse expansion vectors (Fig. S3). Overall, our structures provide many clear routes to developing a range of different potent inhibitors and chemical tool for studying molecular mechanisms of hnRNP mediated splicing regulation.

## Data availability

Atomic coordinates and structure files for the UP1-fragment crystal structures have been deposited in the Protein Data Bank (http://www.pdb.org/). See Table S1 for access codes. Other data are available from the corresponding author upon reasonable request.

## Acknowledgements

F.P. acknowledge support from BBSRC grant (BB/X018997/1) and The Leverhulme Trust grant (RPG-2018-230). This work was supported by funds from the National Institute of General Medical Sciences with grant no. R35GM133488 (to V.V.). Data acquisition was carried out as part of the XCHEM project LB18944-1. We thank the Diamond XCHEM team for their support and expertise.

