## Supporting Information for "Enhanced identification of small molecules binding to hnRNPA1 via cryptic pockets mapping coupled with X-Ray fragment screening"

**Table S1 – Summary of UP1 deposited structures**. The table summarises all deposited UP1-fragment complexes structures. Fragments Z106579662 and Z235361315 binds UP1 on two different surfaces, while two alternative conformations are visible for Z641230552.

| PDB ID | Ligand ID | SMILE | Ligand name |
|---|---|---|---|
| 9F7H | Z45617795 | CS(=O)(=O)NCCC1=CC=CC=C1 | N-(2-phenylethyl)methanesulfonamide |
| 9F5G | EN300-115958 | CS(=O)(=O)N1CCC[C@@H]1CN | [(2R)-1-methanesulfonylpyrrolidin-2-yl]methanamine |
| 9F7F | Z1491353358 | C1CN(C[C@H]1N)S(=O)(=O)C2=CC=CC=C2.Cl | (3S)-1-(benzenesulfonyl)pyrrolidin-3-amine hydrochloride |
| 9F4T | EN300-805013 | COC1=CC=C(C=C1)NC(=O)CN | 2-amino-N-(4-methoxyphenyl)acetamide |
| 9F51 | Z608065044 | CN(C)S(=O)(=O)C1=CNN=C1 | N,N-dimethyl-1H-pyrazole-4-sulfonamide |
| 9F5K | Z33546965 | COC1=CC=C(C=C1)C(=O)NCC(=O)N | N-(2-amino-2-oxoethyl)-4-methoxybenzamide |
| 9F55 | Z235361315 | CCC(=O)NC1=CC=CC(=C1C)N | N-(3-amino-2-methylphenyl)propanamide |
| 9F4V | EN300-118084 | CN1C=C(C(=N1)C(F)F)C(=O)N | 3-(difluoromethyl)-1-methyl-1H-pyrazole-4-carboxamide |
| 9F5C | Z198195770 | CC(=O)NC1=CC(=CC=C1)NC(=O)N | N-[3-(carbamoylamino)phenyl]acetamide |
| 9F4L | Z104584152 | CN(C)C(=O)CNC1=CC=CC=C1 | 2-Anilino-N,N-dimethylacetamide |
| 9F4Z | Z57040482 | CC(=O)NC1=CC=CC=C1O | 2-Acetamidophenol |
| 9F53 | Z237527902 | CC1=CC(=NN1)NC(=O)C | N-(3-methyl-1H-pyrazol-5-yl)acetamide |
| 9F54 | EN300-197154 | CN1C=C(C(=N1)C2CCCNC2)C(=O)N.Cl | 1-methyl-3-(piperidin-3-yl)-1H-pyrazole-4-carboxamide hydrochloride |
| 9F4D | Z1203107138 | CCN1C=C(C=N1)C(=O)NCC2=CC=C(C=C2)F | 1-ethyl-N-[(4-fluorophenyl)methyl]pyrazole-4-carboxamide |
| 9F4K | ZINC72259689 | Cc1nn(C)cc1C(=O)NCC1CCC1 | N-(cyclobutylmethyl)-1,3-dimethylpyrazole-4-carboxamide |
| 9HQL | Z1152242726 | COC1=NC=C(C=C1)NC(=O)NC2=CC=CS2 | N-(6-methoxypyridin-3-yl)-N'-thiophen-2-ylurea |
| 9HQ9 | Z1401276297 | C1CN(CC12CCOC2)C3=NC=CN=C3 | 7-(pyrazin-2-yl)-2-oxa-7-azaspiro[4.4]nonane |
| 9F4G | Z1373445602 | CN1CCN(CC1=O)C2=C(C=CC=N2)F | 4-(3-Fluoropyridin-2-yl)-1-methylpiperazin-2-one |
| 9F4N | Z137811222 | CC(=O)N1CCN(CC1)CCCC2=CC=CC=C2 | 1-[4-(3-phenylpropyl)piperazin-1-yl]ethan-1-one |
| 9F4O | Z991506900 | CC1CN(CCN1)C2=CC=CC(=N2)C | 3-Methyl-1-(6-methylpyridin-2-yl)piperazine |
| 9F4P | Z1217960891 | CN1CCC(C1=O)OC2=CC(=CC=C2)F | 3-(3-Fluorophenoxy)-1-methylpyrrolidin-2-one |
| 9F4S | Z56880342 | CCNC(=O)NC1=NOC(=C1)C | 3-ethyl-1-(5-methyl-1,2-oxazol-3-yl)urea |
| 9F4U | Z111529496 | COCC(=O)NCC1=NC2=CC=CC=C2N1 | N-(1H-benzimidazol-2-ylmethyl)-2-methoxyacetamide |
| 9F4W | Z54508609 | CC(C)C1=CC=C(C=C1)CN2CCC(CC2)O | 1-{[4-(Propan-2-yl)phenyl]methyl}piperidin-4-ol |
| 9F4X | Z1220452176 | CC(=O)NCCC1=CNC2=C1C=C(C=C2)F | N-[2-(5-fluoro-1H-indol-3-yl)ethyl]acetamide |
| 9F4Y | Z90120418 | COC1=CC(=CC(=C1OC)OC)CN.Cl | 3,4,5-Trimethoxybenzylamine hydrochloride |
| 9F5E | Z30820160 | CCC(=O)NC1=NC(=CS1)C | N-(4-methyl-1,3-thiazol-2-yl)propanamide |
| 9F4H | Z106579662 | CC(C)N1CCN(CC1)C(=O)C2=CC=CS2 | 1-(propan-2-yl)-4-(thiophene-2-carbonyl)piperazine |
| 9F4J | Z416341642 | CC1CN(CCO1)S(=O)(=O)N(C)C(C)C | N,2-dimethyl-N-propan-2-ylmorpholine-4-sulfonamide |
| 9F5D | Z641230552 | C1=CC=C(C=C1)CNC2=CNC(=O)NC2=O | 5-(Benzylamino)pyrimidine-2,4(1h,3h)-dione |
| 9F5F | Z992569480 | CNC1CCCN(C1)C2=NN=CC=C2 | N-methyl-1-(pyridazin-3-yl)piperidin-3-amine |
| 9F50 | Z906021418 | CC(C)C1=NC=C(C(=N1)C(=O)N)Cl | 5-Chloro-2-(propan-2-yl)pyrimidine-4-carboxamide |
| 9F4Q | Z641239276 | C1=CC=C(C=C1)CCNC2=CNC(=O)NC2=O | 5-(Phenethylamino)uracil |
| 9F52 | Z734147462 | CNC1(CCOCC1)CO | [4-(methylamino)oxan-4-yl]methanol |
| 9HQJ | Z86417414 | C1=CC=C(C(=C1)C(=O)NC2=NN=CS2)F | 2-fluoro-N-(1,3,4-thiadiazol-2-yl)benzamide |
| 9F4R | Z802821712 | CCNC1=NN=C(C=C1)C | N-ethyl-6-methylpyridazin-3-amine |

**Table S2** – **Crystallographic and refinement statistics** – Statistics for the highest-resolution bin of reflections are in parentheses.

| Ligand ID | Space group | Cell dimensions (Å, ˚) | Resolution (Å) | Total unique reflections | Completeness (%) | $CC_{1/2}$ | Multiplicity | $R_{merge}$ | Mean <I/sI> | $R_{work}/R_{free}$ | Average B-factor (Å²) | Ligand occupancy | PDB ID |
|---|---|---|---|---|---|---|---|---|---|---|---|---|---|
| Z45617795 | P 1 $2_1$ 1 | 37.957 43.907 56.19 90 94.38 90 | 37.85-1.43 (1.45-1.43) | 33868 (1694) | 98.9 (97.4) | 0.992 (0.816) | 1.77 | 0.055 | 7.2 | 0.179/0.200 | 17.67 | 0.64 | 9F7H |
| EN300-115958 | P 1 $2_1$ 1 | 37.959 43.962 56.041 90 94.33 90 | 55.88-1.80 (1.84-1.80) | 17226 (1023) | 99.9 (100) | 0.983 (0.936) | 1.89 | 0.055 | 9.2 | 0.169/0.203 | 15.45 | 0.83 | 9F5G |
| Z1491353358 | P 1 $2_1$ 1 | 38.06 44.41 55.6 90 94.21 90 | 37.96-1.55 (1.58-1.55) | 26646 (1279) | 98.6 (96.6) | 0.995 (0.718) | 1.79 | 0.048 | 10.9 | 0.187/0.221 | 17.33 | 0.72 | 9F7F |
| EN300-805013 | P 1 $2_1$ 1 | 37.979 43.838 56.109 90 94.18 90 | 37.88-1.42 (1.44-1.42) | 58854 (2122) | 98.6 (93.4) | 0.996 (0.668) | 1.70 | 0.052 | 6.2 | 0.189/0.211 | 18.36 | 0.7 | 9F4T |
| Z608065044 | P 1 $2_1$ 1 | 37.977 43.863 56.08 90 94.28 90 | 55.92-1.50 (1.53-1.50) | 29447 (1450) | 99.3 (96.9) | 0.998 (0.981) | 1.76 | 0.024 | 18.3 | 0.172/0.191 | 16.45 | 0.47 | 9F51 |
| Z33546965 | P 1 $2_1$ 1 | 37.941 43.853 56.082 90 94.34 90 | 55.92-1.70 (1.73-1.70) | 20367 (1059) | 99.9 (99.6) | 0.996 (0.876) | 1.89 | 0.046 | 10.8 | 0.172/0.187 | 17.71 | 0.61 | 9F5K |
| Z235361315 | P 1 $2_1$ 1 | 38.005 43.915 55.917 90 94.05 90 | 37.91-1.55 (1.58-1.55) | 26686 (1245) | 99.3 (93.6) | 0.999 (0.963) | 1.80 | 0.024 | 16.1 | 0.175/0.201 | 17.83 | 0.68/0.64 | 9F55 |
| EN300-118084 | P 1 $2_1$ 1 | 37.999 43.804 55.982 90 94.06 90 | 37.90-1.40 (1.42-1.40) | 35493 (1483) | 97.7 (82.3) | 0.996 (0.879) | 1.72 | 0.039 | 9.4 | 0.181/0.200 | 18.67 | 0.47 | 9F4V |
| Z198195770 | P 1 $2_1$ 1 | 37.999 43.957 56.03 90 94.51 90 | 55.86-1.45 (1.47-1.45) | 32278 (1448) | 98.4 (91.7) | 0.999 (0.957) | 1.79 | 0.026 | 14.5 | 0.175/0.208 | 16.83 | 0.82 | 9F5C |
| Z104584152 | P 1 $2_1$ 1 | 38.181 43.929 55.749 90 93.56 90 | 55.64-1.40 (1.42-1.40) | 59339 (1959) | 97.6 (81.1) | 0.998 (0.851) | 1.64 | 0.030 | 10.1 | 0.183/0.204 | 19.19 | 0.8 | 9F4L |
| Z57040482 | P 1 $2_1$ 1 | 38.02 43.959 55.922 90 94.23 90 | 37.92-1.60 (1.63-1.60) | 24436 (1204) | 99.9 (92.8) | 0.996 (0.928) | 1.86 | 0.040 | 10.1 | 0.172/0.203 | 19.75 | 0.78 | 9F4Z |
| Z237527902 | P 1 $2_1$ 1 | 38.1 43.968 55.948 90 94.2 90 | 34.53-1.80 (1.84-1.80) | 16317 (959) | 95.2 (94.4) | 0.997 (0.955) | 1.68 | 0.032 | 19.9 | 0.162/0.191 | 16.51 | 0.74 | 9F53 |
| EN300-197154 | P 1 $2_1$ 1 | 38.065 43.889 56.104 90 94.19 90 | 37.96-1.70 (1.73-1.70) | 20474 (1064) | 99.9 (99.5) | 0.998 (0.937) | 1.87 | 0.033 | 13.9 | 0.171/0.197 | 18.85 | 0.58 | 9F54 |
| Z1203107138 | P 1 $2_1$ 1 | 38.05 43.928 55.955 90.00 94.01 90.00 | 37.96-1.50 (1.53-1.50) | 29579 (1484) | 99.7 (99.5) | 0.999 (0.901) | 1.86 | 0.023 | 19.0 | 0.175/0.194 | 17.86 | 0.39 | 9F4D |
| ZINC72259689 | P 1 $2_1$ 1 | 38.29 44.05 56.15 90 93.99 90 | 56.01-1.80 (1.84-1.80) | 17394 (1038) | 99.6 (99.6) | 0.998 (0.934) | 1.87 | 0.034 | 19.5 | 0.173/0.220 | 20.69 | 0.93 | 9F4K |
| Z1152242726 | P 1 $2_1$ 1 | 37.90 43.92 55.73 90 94.05 90 | 55.59-1.75 (1.78-1.75) | 17906 (950) | 96.6 (95.8) | 0.981 (0.767) | 1.80 | 0.092 | 11.1 | 0.189/0.212 | 21.17 | 0.65 | 9HQL |
| Z1401276297 | P 1 $2_1$ 1 | 37.00 44.09 55.98 90 94.17 90 | 55.84-1.60 (1.63-1.60) | 24516 (1218) | 99.8 (99.6) | 0.998 (0.741) | 1.83 | 0.050 | 11.33 | 0.179/0.211 | 16.92 | 0.71 | 9HQ9 |
| Z1373445602 | P 1 $2_1$ 1 | 37.943 43.855 55.829 90 94.12 90 | 37.84-1.40 (1.42-1.40) | 61143 (2787) | 99.6 (99.3) | 0.999 (0.579) | 1.71 | 0.035 | 12.7 | 0.186/0.221 | 17.28 | 0.49 | 9F4G |
| Z137811222 | P 1 $2_1$ 1 | 38.054 43.909 56.043 90 94.1 90 | 37.96-1.40 (1.42-1.40) | 61276 (2051) | 98.4 (85.7) | 0.987 (0.949) | 1.70 | 0.051 | 10.1 | 0.180/0.197 | 16.82 | 0.57 | 9F4N |
| Z991506900 | P 1 $2_1$ 1 | 37.864 43.968 56.083 90 94.64 90 | 37.74-1.40 (1.41-1.40) | 35631 (1558) | 98.1 (88.4) | 0.992 (0.969) | 1.80 | 0.043 | 11.9 | 0.168/0.181 | 14.68 | 0.84 | 9F4O |
| Z1217960891 | P 1 $2_1$ 1 | 38.02 43.868 56.094 90 94.16 90 | 55.95-1.40 (1.42-1.40) | 35810 (1525) | 98.3 (84.4) | 0.995 (0.909) | 1.67 | 0.039 | 9.9 | 0.189/0.205 | 17.72 | 0.61 | 9F4P |
| Z56880342 | P 1 $2_1$ 1 | 38.039 43.655 56.065 90 94.05 90 | 34.41-1.40 (1.42-1.40) | 35627 (1512) | 98.3 (84.9) | 0.988 (0.946) | 1.72 | 0.054 | 9.0 | 0.178/0.189 | 16.66 | 0.7 | 9F4S |
| Z111529496 | P 1 $2_1$ 1 | 38.029 43.907 56.007 90 94.14 90 | 37.93-1.40 (1.42-1.40) | 35599 (1541) | 97.8 (85.8) | 0.994 (0.951) | 1.69 | 0.045 | 9.8 | 0.175/0.193 | 17.46 | 0.5 | 9F4U |
| Z54508609 | P 1 $2_1$ 1 | 37.955 43.807 56.143 90 94.46 90 | 37.84-1.60 (1.63-1.60) | 24387 (1214) | 99.8 (99.8) | 0.987 (0.912) | 1.85 | 0.062 | 7.6 | 0.178/0.209 | 17.11 | 0.68 | 9F4W |
| Z1220452176 | P 1 $2_1$ 1 | 38.063 43.855 55.951 90 93.9 90 | 37.97-1.60 (1.63-1.60) | 24400 (1201) | 99.8 (99.8) | 0.996 (0.939) | 1.83 | 0.042 | 12.0 | 0.179/0.207 | 16.46 | 0.63 | 9F4X |
| Z90120418 | P 1 $2_1$ 1 | 38.029 43.882 55.875 90 94.19 90 | 37.93-1.40 (1.42-1.40) | 61536 (2026) | 98.1 (84.6) | 0.999 (0.969) | 1.71 | 0.019 | 16.5 | 0.173/0.190 | 17.39 | 0.74 | 9F4Y |
| Z30820160 | P 1 $2_1$ 1 | 37.893 43.977 56.013 90 94.56 90 | 37.77-1.45 (1.47-1.45) | 32400 (1490) | 99.1 (95.0) | 0.997 (0.966) | 1.72 | 0.029 | 12.8 | 0.170/0.189 | 15.72 | 0.66 | 9F5E |
| Z106579662 | P 1 $2_1$ 1 | 38.062 43.938 55.928 90 94.38 90 | 37.95-1.40 (1.42-1.40) | 34576 (1383) | 95.2 (76.7) | 0.998 (0.742) | 1.71 | 0.036 | 10.3 | 0.180/0.205 | 17.89 | 0.54/0.68 | 9F4H |
| Z416341642 | P 1 $2_1$ 1 | 38.006 43.9 56.009 90 94.22 90 | 34.52-1.40 (1.42-1.40) | 61846 (2099) | 96.8 (90.9) | 0.998 (0.704) | 1.72 | 0.032 | 10.5 | 0.182/0.198 | 20.02 | 0.64 | 9F4J |
| Z641230552 | P 1 $2_1$ 1 | 38.022 43.796 55.799 90 93.34 90 | 37.96-1.60 (1.63-1.60) | 23932 (1156) | 98.5 (98.2) | 1.000 (0.970) | 1.79 | 0.017 | 19.9 | 0.176/0.199 | 21.31 | 0.66/0.34 | 9F5D |
| Z992569480 | P 1 $2_1$ 1 | 37.977 43.898 55.937 90 94.19 90 | 32.46-1.40 (1.42-1.40) | 35726 (1525) | 98.4 (85.4) | 0.986 (0.913) | 1.73 | 0.058 | 7.8 | 0.178/0.204 | 16.42 | 0.71 | 9F5F |
| Z906021418 | P 1 $2_1$ 1 | 38.019 43.907 56.168 90 94.36 90 | 37.91-1.50 (1.53-1.50) | 29582 (1445) | 99.4 (97.4) | 0.998 (0.973) | 1.80 | 0.026 | 16.8 | 0.174/0.192 | 17.7 | 0.68 | 9F50 |
| Z641239276 | P 1 $2_1$ 1 | 37.93 43.934 55.973 90 94.07 90 | 55.83-1.40 (1.42-1.40) | 54783 (1418) | 92.0 (65.4) | 0.995 (0.898) | 1.52 | 0.043 | 9.1 | 0.182/0.201 | 18.82 | 0.77 | 9F4Q |
| Z734147462 | P 1 $2_1$ 1 | 38.056 44.029 56.055 90 94.38 90 | 37.95-1.50 (1.53-1.50) | 29582 (1417) | 99.3 (94.8) | 1.000 (0.985) | 1.81 | 0.014 | 23.9 | 0.174/0.195 | 17.45 | 0.56 | 9F52 |
| Z86417414 | P 1 $2_1$ 1 | 37.944 43.778 55.839 90 94.027 90 | 55.70-1.60 (1.63-1.60) | 24249 (1185) | 99.8 (99.9) | 0.998 (0.719) | 1.89 | 0.045 | 12.3 | 0.182/0.210 | 18.23 | 0.64 | 9HQJ |
| Z802821712 | P 1 $2_1$ 1 | 38.106 44.045 56.217 90 94.23 90 | 56.06-1.59 (1.62-1.59) | 25086 (1203) | 99.3 (94.6) | 0.981 (0.797) | 2.81 | 0.089 | 8.9 | 0.186/0.217 | 22.07 | 0.86 | 9F4R |

**Table S3 – Identified fragments**. The table summarises information regarding fragments identified to bind UP1, highlighting fragment induced changes in F17 and F59 side chains and inter-RRM loop conformations. Furthermore, fragment binding locations are indicated.

| Ligand ID | F17 and F59 conformations | Inter-RRM loop structure | Binding site(s) | Figure 1 relative binding sites |
|---|---|---|---|---|
| Z45617795 | alternative conformations | flexible | RRM1<br>α1 - (α2-β4) loop | 5 |
| EN300-115958 | alternative conformations | flexible | RRM1<br>α1 - (α2-β4) loop | 5 |
| Z1491353358 | RNA-bound conformation | flexible | S2 | |
| EN300-805013 | alternative conformations | flexible | inter-RRMs (α2-α2) | 4 |
| Z608065044 | alternative conformations | flexible | inter-RRMs (α2-α2) | 4 |
| Z33546965 | alternative conformations | flexible | inter-RRMs (α2-α4) | 4 |
| Z235361315 | alternative conformations | flexible | inter-RRMs (α2-α2); S1 | 4 |
| EN300-118084 | alternative conformations | flexible | S3 | |
| Z198195770 | alternative conformations | flexible | S3 | |
| Z104584152 | alternative conformations | flexible | S1 | |
| Z57040482 | alternative conformations | rigid | inter-RRMs loop | 3 |
| Z237527902 | alternative conformations | rigid | inter-RRMs loop | 3 |
| EN300-197154 | alternative conformations | flexible | RRM2 (β2-β3) loop | 6 |
| Z1203107138 | alternative conformations | flexible | nucleobase pocket 1 | 1 |
| ZINC72259689 | F17 in alternative conformation;<br>F59 anti-RNA-bound conformation | flexible | nucleobase pocket 1 | 1 |
| Z137811222 | F17 in alternative conformation;<br>F59 anti-RNA-bound conformation | flexible | nucleobase pocket 1 | 1 |
| Z1217960891 | F17 in alternative conformation;<br>F59 anti-RNA-bound conformation | flexible | nucleobase pocket 1 | 1 |
| Z1373445602 | F17 RNA-bound conformation<br>(F17B very low occupancy/not fully visible);<br>F59 alternative conformation | flexible | nucleobase pocket 1 | 1 |
| Z1401276297 | F17 RNA-bound conformation;<br>F59 alternative conformation | flexible | nucleobase pocket 1 | 1 |
| Z106579662 | F17 RNA-bound conformation<br>(F17B very low occupancy/not fully visible);<br>F59 alternative conformation | flexible | nucleobase pocket 1; S1 | 1 |
| Z991506900 | RNA-bound conformation | rigid | nucleobase pocket 1 | 1 |
| Z30820160 | RNA-bound conformation;<br>(F59B very low occupancy/not fully visible) | rigid | nucleobase pocket 1 | 1 |
| Z56880342 | anti-RNA-bound conformation | flexible | nucleobase pocket 1 | 1 |
| Z54508609 | anti-RNA-bound conformation | flexible | nucleobase pocket 1 | 1 |
| Z1220452176 | anti-RNA-bound conformation | flexible | nucleobase pocket 1 | 1 |
| Z90120418 | anti-RNA-bound conformation | flexible | nucleobase pocket 1 | 1 |
| Z1152242726 | anti-RNA-bound conformation | flexible | nucleobase pocket 1 | 1 |
| Z111529496 | alternative conformations | flexible | nucleobase pocket 1 | 1 |
| Z416341642 | alternative conformations | flexible | nucleobase pocket 2 | 2 |
| Z641230552 | alternative conformations | flexible | nucleobase pocket 2 | 2 |
| Z992569480 | alternative conformations | flexible | nucleobase pocket 2 | 2 |
| Z906021418 | alternative conformations | flexible | nucleobase pocket 2 | 2 |
| Z641239276 | F17 RNA-bound conformation;<br>F59 alternative conformation | flexible | S4 | |
| Z734147462 | alternative conformations | flexible | RRM2 α2 - β4 | 7 |
| Z86417414 | alternative conformations | flexible | RRM2 α1 - β2 | 6 |
| Z802821712 | alternative conformations | flexible | RRM2 α1 - β2 | 6 |

S1 – fragment sandwiched between RRM1 loop (β1-β3) and RRM2 α1 of the symmetry related molecule; S2 – fragment sandwiched among 3 symmetry related molecules: inter-RRMs interface, the RRM1 (β2 – β3) loop of one symmetry related molecules and the RRM2 (β2 – β3) loop of the second symmetry related molecule; S3 – fragment sandwiched between (inter-RRMs loop – (α5) loop) and RRM2 (β2- β3)-loop of the symmetry related molecule; S4 - fragment sandwiched between nucleobase pocket 2 and RRM2 α1 – β3 of the symmetry related molecule.

**Table S4 – UP1-fragment interactions**. UP1 residues forming contacts with fragments in the structures are listed by their position and by their single-letter identity. Interactions were identified using PyMol.

| Ligand ID | Binding site(s) | Van der Walls | π-stacking | π-cation | H-bond | Salt bridge | Halogen bond | Water bridge |
|---|---|---|---|---|---|---|---|---|
| Z45617795 | RRM1 α1 - (α2-β4) loop | P76, V83 | | | K78, P76 | | | A74 |
| EN300-115958 | RRM1 α1 - (α2-β4) loop | | | | K78 | | | P76 |
| Z1491353358 | S2 | L13, I164, T26(1*) | | | N50(2*), Q12, E24(1*) | | | |
| EN300-805013 | inter-RRMs (α2-α2) | | | K161 | N73, D69 | | | R75, Q127, K161 |
| Z608065044 | inter-RRMs (α2-α2) | | | | E126, K130, K161 | | | N73, R75, D157 |
| Z33546965 | inter-RRMs (α2-α2) | | | | N73 | | | D69, R75, Q127, K161 |
| Z235361315 | inter-RRMs (α2-α2) | | | | Q127 | | | E126, G129, K161 |
| | S1 | V65(1*) | | | T51, E66(1*) | | | R47 |
| EN300-118084 | S3 | | | | K105, H156 | D155, D139(1*) | T103 | S142(1*) |
| Z198195770 | S3 | | | | K105, D155, H156, D139(1*) | | | R146(1*) |
| Z104584152 | S1 | V65 | | | R47(1*) | E66 | | F23(1*) |
| Z57040482 | inter-RRMs loop | L102, V163, I164 | | | S91, R92, L102, K179 | | | |
| Z237527902 | inter-RRMs loop | L102, I164 | | | S91, R92, L102, K179 | | | |
| EN300-197154 | RRM2 (β2-β3) loop | | | | T138, D139, R140 | | | |
| Z1203107138 | nucleobase pocket 1 | F17, H101 | F57 | | | | R55 | |
| ZINC72259689 | nucleobase pocket 1 | F17, F57, F59, A89 | F17 | | V90 | | | |
| Z137811222 | nucleobase pocket 1 | F57, F59, A89 | | | | | | |
| Z1217960891 | nucleobase pocket 1 | F17, F57, F59 | | | | | | |
| Z1373445602 | nucleobase pocket 1 | F17, F57 | | | | | | K87 |
| Z1401276297 | nucleobase pocket 1 | F59 | F17 | | R55 | | | |
| Z106579662 | nucleobase pocket 1 | F57, A90 | | | H101 | | | K87 |
| | S1 | V65 | | | R47(1*) | E9, E66 | | S54(1*), F23(1*) |
| Z991506900 | nucleobase pocket 1 | F17, F57, F59, A89 | | | | | | K87 |
| Z30820160 | nucleobase pocket 1 | F17, A89 | | | | | | K87, G99 |
| Z56880342 | nucleobase pocket 1 | F17, F57, F59 | | | | | | E85 |
| Z54508609 | nucleobase pocket 1 | F17, F57, F59 | F17, F57 | | | | | G20, G56 |
| Z1220452176 | nucleobase pocket 1 | F17, F57, F59 | | | | | | |
| Z90120418 | nucleobase pocket 1 | F17, F57 | | | | | | V90, H101 |
| Z111529496 | nucleobase pocket 1 | F17, F57, F59 | | | | | | |
| Z1152242726 | nucleobase pocket 1 | | F17 | | V90 | | | K87, R88 |
| Z416341642 | nucleobase pocket 2 | F108, F148, A180 | | | R178 | | | |
| Z641230552 | nucleobase pocket 2 | A180, F150 | | | K106, E135, L181 | | | |
| Z992569480 | nucleobase pocket 2 | F108, F148 | | | | | | L181 |
| Z906021418 | nucleobase pocket 2 | F108 | F108 | | | | | L181 |
| Z641239276 | S4 | F150, I131(1*), I134(1*) | | | K106,E135, K130(1*) | | | |
| Z734147462 | RRM2 α2 - β4 | | | | Q165, V177 | | | K166 |
| Z86417414 | RRM2 α1 - β2 | I131, I134 | | | E118, I134 | | E118, R122 | |
| Z802821712 | RRM2 α1 - β2 | I131 | | | R122, I134 | | | I134 |

(n*) = symmetry related

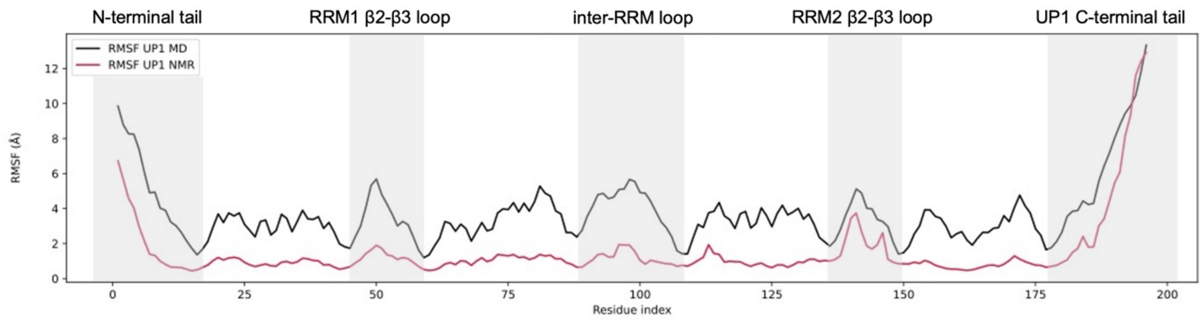


**Figure S1 Comparison of MD simulations and NMR data**. Per-residue root mean square fluctuation (RMSF) for the extended MD simulation (black line) was superimposed for direct comparison over the RMSF of the NMR ensemble (PDB ID: 2LYV) (red line). Flexible regions are highlighted with grey boxes.

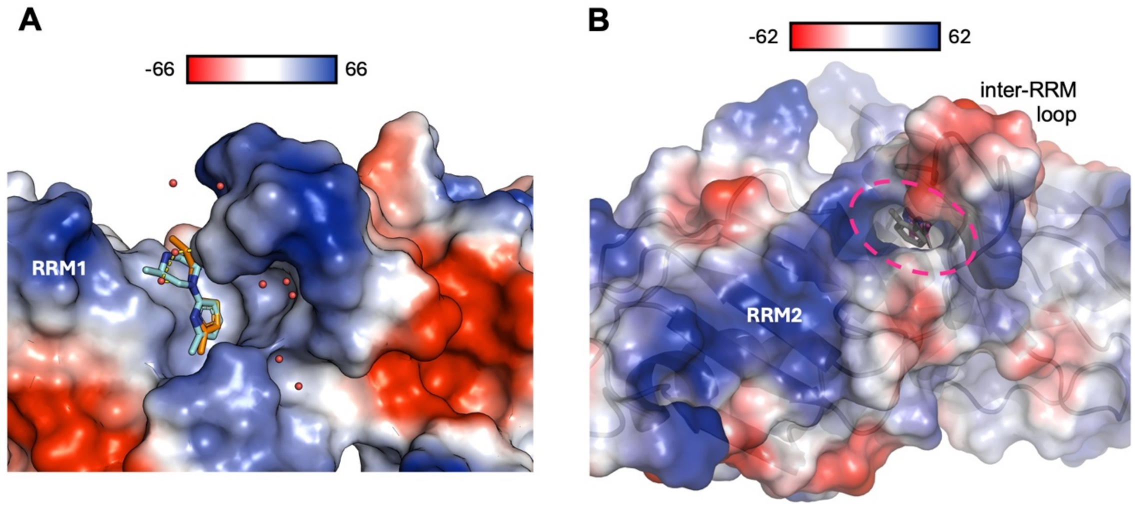


**Figure S2 Fragments induced conformational changes**. **A**) UP1 electrostatic surface showing the formation of a positive charged pocket at the edge on RRM1 upon binding of Z30820160 or Z991506900 to the nucleobase pocket 1. **B**) UP1 electrostatic surface showing the formation of a mostly neutral pocket (highlighted with a magenta dotted line) on the edge of RRM2 upon binding of Z57040482 and Z237527902 to the inter-RRM loop site.

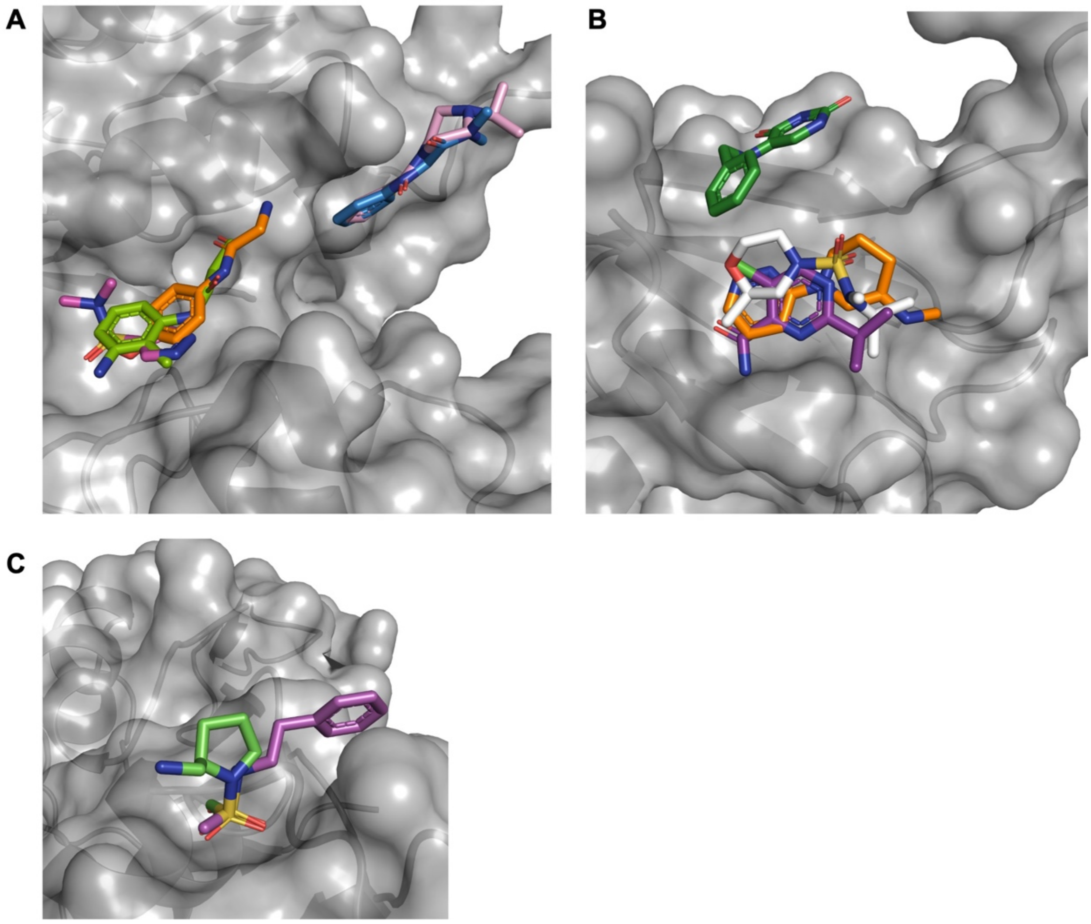


**Figure S3. Fragment merging and linking opportunities can be directly inferred for many hits. A**) Overlay of EN300-805013 (orange), Z608065044 (magenta), Z235361315 (green), Z104584152 (blue) and Z106579662 (pink). **B**) Z416341642 (white), Z992569480 (orange), Z906021418 (purple), Z641230552 (green). **C**) Overlay of Z45617795 (magenta) and EN300-115958 (green).

**A**

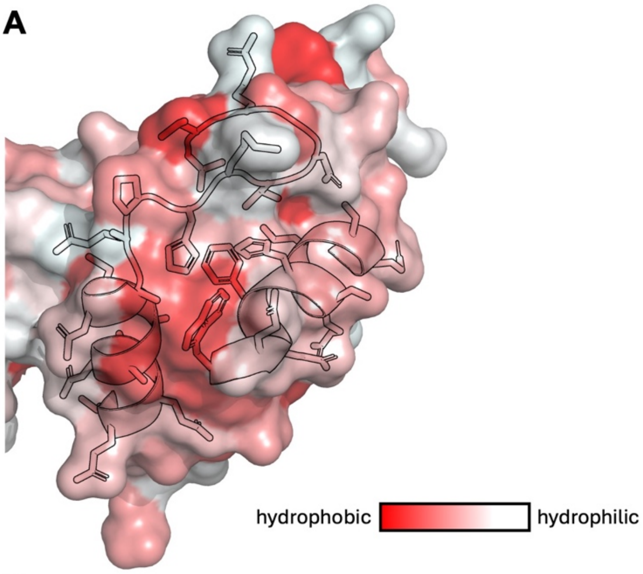


**B**

```
                β1          α1         β2           β3        α2
UP1_RRM1        --RKLFIGGLSFETTDESLRSHFEQWGTLTDCVVMRDPNTKRSRGFGFVTYATVEEVDAA
UP1_RRM2        --KKIFVGGIKEDTEEHHLRDYFEQYGKIEVIEIMTDRGSGKKRGFAFVTFDDHDSVDKI
hnRNPL_RRM2     SVLLFTILNPIYSITTDVLYTICNPCGPVQRIVIFRK-NG----VQAMVEFDSVQSAQRA
FIR_RRM2        --NRIYVASVHQDLSDDDIKSVFEAFGKIKSCTLARDPTTGKHKGYGFIEYEKAQSSQDA
PTB_RRM1        --RVIHIRKLPIDVTEGEVISLGLPFGKVTNLLMLKGKNQ------AFIEMNTEEAANTM
PTB_RRM2        --LRIIVENLFYPVTLDVLHQIFSKFGTVLKIITFTKNNQ----FQALLQYADPVSAQHA
CBP20           --CTLYVGNLSFYTTEEQIYELFSKSGDIKKIIMGLDKMKKTACGFCFVEYYSRADAENA
RBM7            --RTLFVGNLETKVTEELLFELFHQAGPVIKVKIPKDKDG-KPKQFAFVNFKHEVSVPYA

                         β4
UP1_RRM1        MNARP---HKVDGRVVEPKRAVSREDSQR----
UP1_RRM2        VIQKY---HTVNGHNCEVRKALSKQ--------
hnRNPL_RRM2     KASLNG--ADIYSGCCTLKIEYAKP--------
FIR_RRM2        VSSMNL--FDLGGQYLRVGKAVT----------
PTB_RRM1        VNYYTSVTPVLRGQPIYIQFSNHKELKTDSSPN
PTB_RRM2        KLSLDG--QNIYNACCTLRIDFSKL--------
CBP20           MRYING--TRLDDRIIRTDWDAG----------
RBM7            MNLLNG--IKLYGRPIKIQFRSG----------
```

**Figure S4. RRM PPI surface. A**) Conserved hydrophobic depression on the predicted PPI surface defined by RRM1 α1 - α2 – (α2-β4) loop of UP1. UP1 RRM1 surface is coloured according to the Eisenberg hydrophobicity scale. **B**) Sequence alignment among the RRM1 and 2 of UP1 and the RRM domains of hnRNPL, FBP-interacting repressor (FIR), Polypyrimidine tract binding protein (PTB), Nuclear cap-binding protein subunit 2 (CBP20), RNA-binding protein 7 (RBM7) sequences, mapped against the conserved RRM secondary structure. In bold are highlighted residues mediating fragments-UP1 interactions in our structures and protein-protein/peptide interactions in hnRNPL, FIR, PTB RRM1 and RRM2, CBP20 and RBM7 structures (PDB ID 7EVR, 2KXH, 8BWF, 3ZZY, 5oo6, 5LXY, respectively).

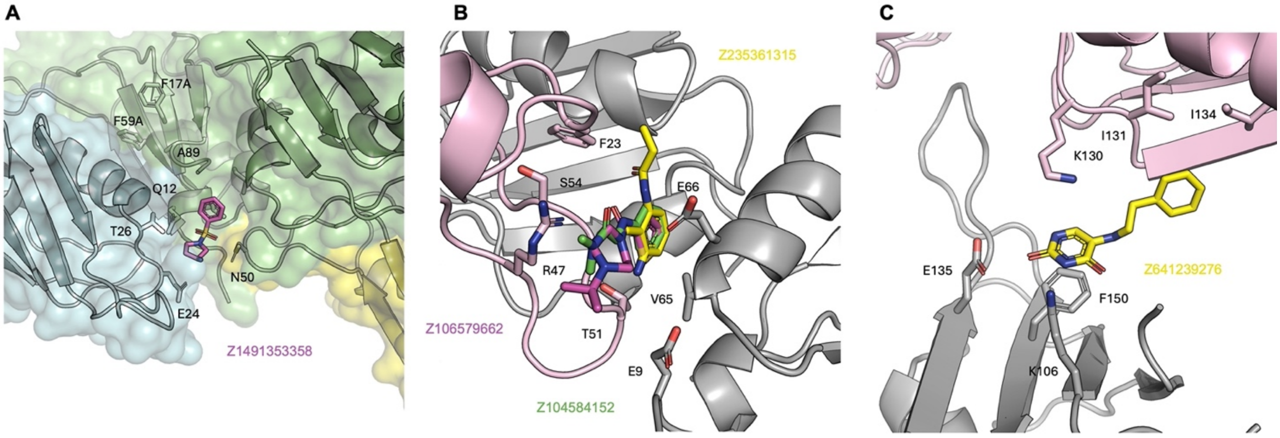


**Figure S5 Fragments Influenced by Crystal Contacts**. **A**) Z1491353358 (purple) bound on a pocket generated at the intersection of three symmetry-related molecules, and formed by the inter-RRMs interface of molecules 1 (green), RRM1 (β2 – β3) loop of symmetry related molecules 1 (yellow) and RRM1 (β1 – α1) loop of symmetry related molecule 2 (cyan). **B)** overlay of Z235361315 (yellow), Z104584152 (green), and Z106579662 (purple) bound to the inter-RRMs interface (grey) and the RRM1 (β2 – β3) loop of the symmetry-related molecules (pink). **C**) Z641239276 (yellow) bound to the RRM2 nucleobase pocket (grey) and the RRM2 α1 – β3 of the symmetry-related molecules (pink).
